# COVIDep platform for real-time reporting of vaccine target recommendations for SARS-CoV-2: Description and connections with COVID-19 immune responses and preclinical vaccine trials

**DOI:** 10.1101/2020.05.23.111385

**Authors:** Syed Faraz Ahmed, Ahmed A. Quadeer, Matthew R. McKay

## Abstract

We introduce COVIDep (https://COVIDep.ust.hk), a web-based platform that provides immune target recommendations for guiding SARS-CoV-2 vaccine development. COVIDep implements a protocol that pools together publicly-available genetic data for SARS-CoV-2 and epitope data for SARS-CoV to identify B cell and T cell epitopes that present potential immune targets for SARS-CoV-2. Correspondences between outputs of COVIDep and immune responses recorded in COVID-19 patients and preclinical vaccine trials are also indicated. The platform is user-friendly, flexible, and based on up-to-date data. It may help guide vaccine designs and associated experimental studies for SARS-CoV-2.

## Description

The COVID-19 pandemic, caused by the novel coronavirus SARS-CoV-2, has brought much of the world to a virtual lockdown. As the virus continues to spread rapidly and the pandemic intensifies, the need for an effective vaccine is becoming increasingly apparent. A critical part of vaccine design is to identify targets, or epitopes, that can induce an effective immune response against SARS-CoV-2. This problem is challenged by our limited understanding of this novel coronavirus and of its interplay with the human immune system.

In response to this challenge, we have developed COVIDep (https://COVIDep.ust.hk), a first-of-its-kind web-based platform that pools genetic data for SARS-CoV-2 and immunological data for the 2003 SARS virus, SARS-CoV, to identify B cell and T cell epitopes to serve as vaccine target recommendations for SARS-CoV-2 (Figure 1). For T cell epitopes, it provides estimates of population coverage, globally and for specific regions. The COVIDep platform is updated periodically as data is deposited into public databases. This is important since SARS-CoV-2 sequences are being made available at an increasing rate through international data sharing efforts, and the identification of vaccine targets is influenced by newly observed genetic variation. COVIDep is flexible and user-friendly, comprising an intuitive graphical interface and interactive visualizations.

**Figure 1.**
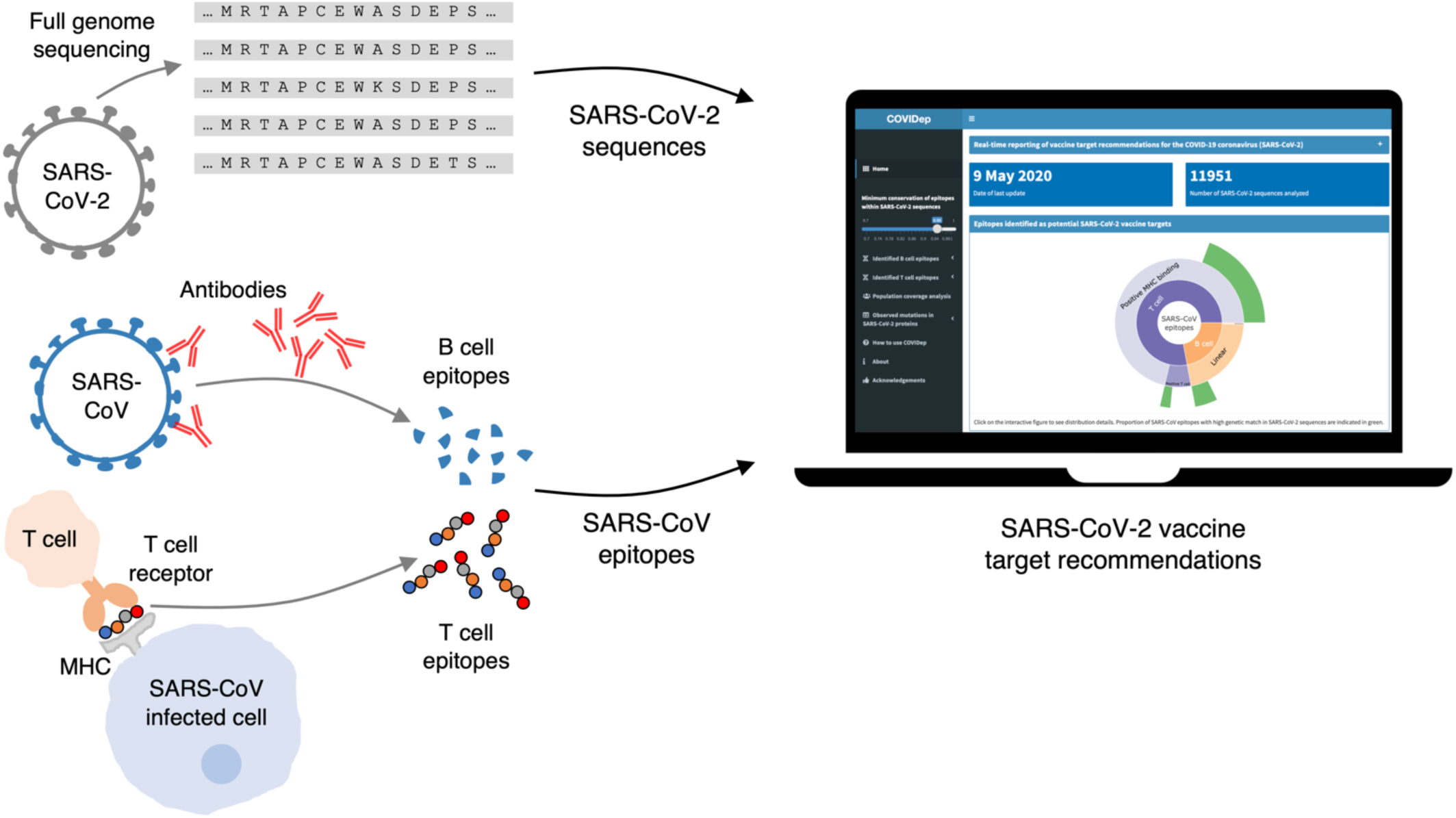
COVIDep provides sets of B cell and T cell epitopes that can serve as potential vaccine targets for SARS-CoV-2. The recommended epitopes are experimentally-derived from SARS-CoV and have a close genetic match with the available SARS-CoV-2 sequences (see Supplementary Figure 1 for a detailed protocol description).

The vaccine targets recommended by COVIDep exploit the genetic similarities between SARS-CoV-2 and SARS-CoV, along with known immune targets for SARS-CoV that have been determined experimentally. The system implements a protocol that identifies from among the SARS epitopes that can induce a human immune response, those that are genetically similar in SARS-CoV-2. This idea, put forward in our preliminary study^1^ based on limited early data, identified known SARS-CoV epitopes that had an identical genetic match in SARS-CoV-2. These epitopes presented initial vaccine target recommendations for potentially eliciting a protective, cross-reactive immune response against SARS-CoV-2. Similar ideas and results were reported subsequently in an independent study^2^.

The use of SARS-CoV immunological data to inform vaccine targets for SARS-CoV-2 is being supported by experimental results. There is evidence of SARS-CoV-derived antibodies binding to genetically similar regions of SARS-CoV-2’s spike protein^3^, and also of cross-neutralization^4–6^. Conversely, studies have demonstrated that specific SARS-CoV-derived antibodies binding to the spike’s receptor binding domain, which has significant genetic differences in SARS-CoV-2, have limited cross-reactivity^7^. Antibody and T cell responses against spike protein epitopes that are genetically similar in SARS-CoV and SARS-CoV-2 have also been reported in COVID-19 infected patients^8–10^, and in preclinical vaccine trials^11,12^. Epitopes recommended by COVIDep have notable overlap with the findings in these experimental studies (Figures 2 and 3).

**Figure 2.**
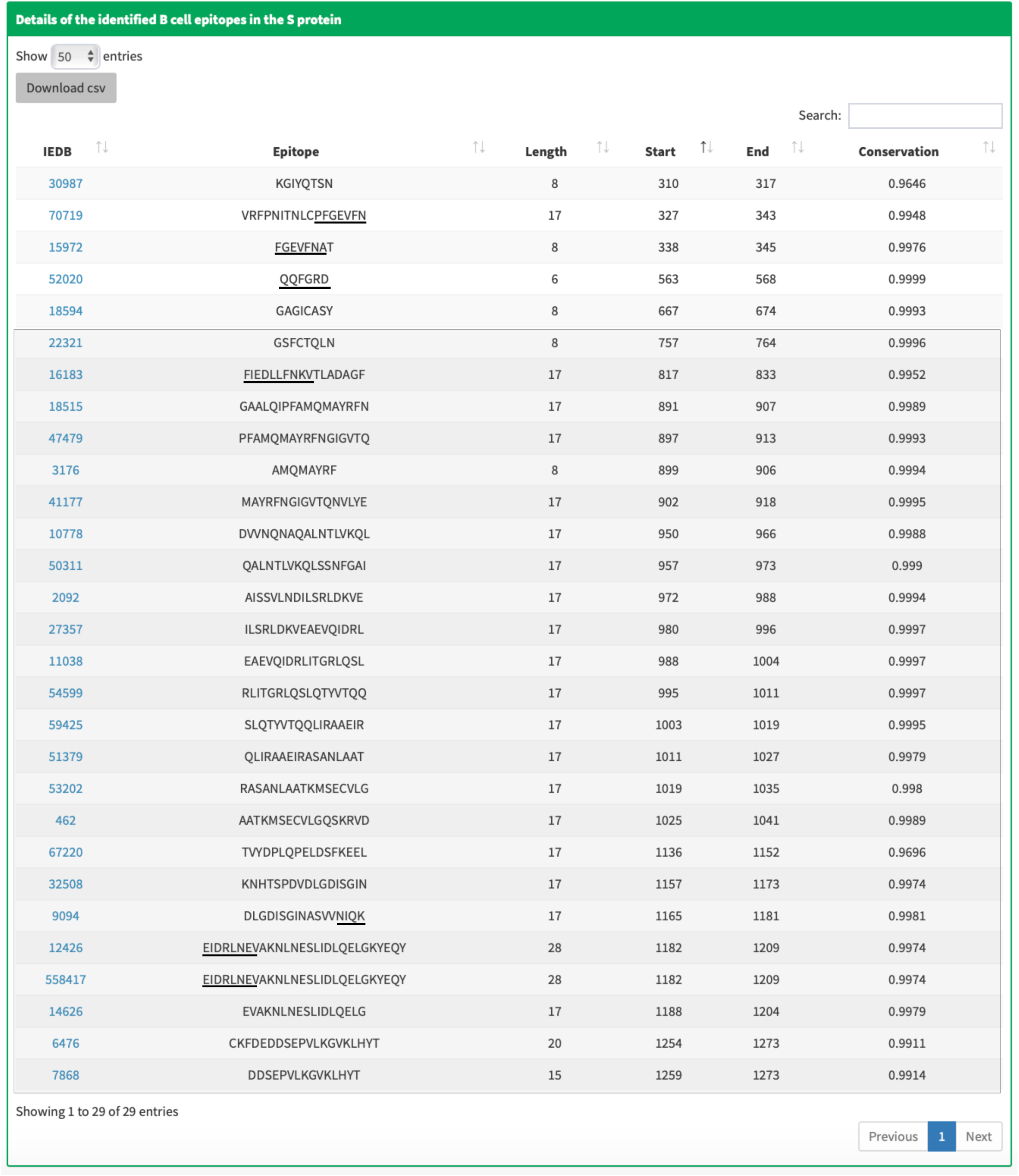
B cell linear epitopes in the spike protein of SARS-CoV-2 identified by COVIDep (as of 21 May 2020) and their overlap with emerging experimental results. The majority of the identified epitopes (24/29; shown in a shaded box) are located in the S2 functional subunit of the spike protein, reported to be a main region targeted by cross-reactive^3^ and cross-neutralizing^4^ antibodies. The epitopes with IEDB IDs 70719 and 15972 overlap with regions in the S1 functional subunit of the spike protein reported to be targeted by cross-neutralizing antibodies^5,6^. The specific overlapping residues are underlined. The epitopes with IEDB IDs 9094, 12426, and 558417 overlap with an epitope (located at positions 1178-1189) reported to be targeted by neutralizing antibodies in a preclinical trial of a SARS-CoV-2 vaccine candidate^11^. Interestingly, the partial overlaps of the (consecutive) epitopes 9094 and 12426/558417 cover the experimentally-reported epitope^11^ completely. Note that epitopes 12426 and 558417 share the same sequence; they have different IDs due to differences in the associated experimental procedures. The epitopes with IEDB IDs 52020 and 16183 overlap with the regions reported to be recognized by neutralizing antibodies in the sera of recovered COVID-19 patients^8^.

**Figure 3.**
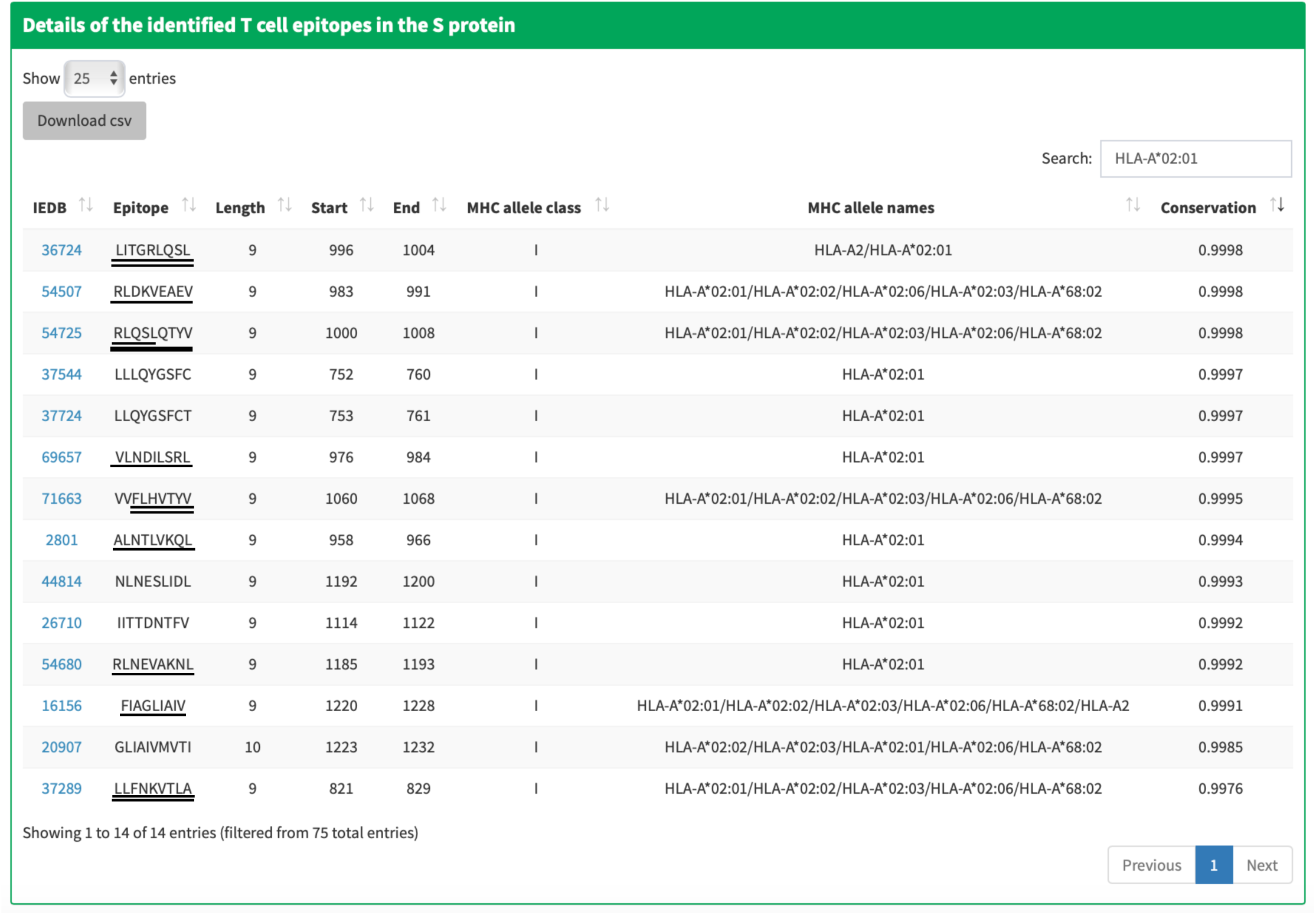
T cell epitopes in the spike protein of SARS-CoV-2 identified by COVIDep (as of 21 May 2020) and their overlap with emerging experimental results. Of the 14 HLA-A*02:01-restricted spike protein epitopes identified by COVIDep, nine epitopes (IEDB IDs 36724, 54507, 54725, 69657, 71663, 2801, 54680, 16156, and 37289) overlap, completely or significantly, with epitopes against which cytotoxic CD8+ T cell responses have been observed in peripheral blood mononuclear cells isolated from COVID-19 patients^9,10^. In a preclinical vaccine trial^12^, T cell responses have also been recorded against a protein region comprising the identified epitope with IEDB ID 71663. The specific overlapping residues are underlined, with double-underline reflecting that a response was observed in two studies. A more prominent underline is used to distinguish the epitope with IEDB ID 54725, since a response against this epitope was observed in ten COVID-19 patients (of fourteen studied)^10^. Therefore, this epitope appears to be particularly immunogenic. The epitopes with IEDB IDs 36724, 69657, 71663, 2801, 54680, and 16156 were originally reported based on positive T cell assays for SARS-CoV, while those with IEDB IDs 54507, 54725, and 37289 were reported based on positive MHC binding assays.

The recommendations provided by COVIDep may be used to guide vaccine designs and associated experimental studies, and may help to expedite the discovery of an effective vaccine for COVID-19.

## Data availability

The SARS-CoV-2 full genome sequence data is periodically downloaded from the Global Initiative on Sharing Avian Influenza Database (GISAID; www.gisaid.org). The SARS-CoV epitope sequence data was downloaded from the Virus Pathogen Database and Analysis Resource (ViPR; www.viprbrc.org). The population coverage statistics of HLA alleles were obtained from the Immune Epitope Database and Analysis Resource (IEDB; www.iedb.org).

## Code availability

The source code for the developed platform is available at the COVIDep GitHub repository (https://github.com/COVIDep).

## Acknowledgment

COVIDep is made possible by the open sharing of genome sequence data of SARS-CoV-2 sequences by research groups from around the world through the GISAID platform, and the open sharing of immunological data of experimentally-determined SARS-CoV epitopes through the ViPR database. We gratefully acknowledge the contributions of all the researchers, scientists and technical staff involved (a detailed acknowledgment is available at the Acknowledgments page of the COVIDep platform).

We thank Nelvin Law and Kenny Pang for their technical support, and Raymond Louie, David Morales-Jimenez, Saqib Sohail, Neelkanth Kundu, Awais Shah, and Umer Abdullah for comments and suggestions that helped to improve the web application and its interface.

M.R.M. and A.A.Q. were supported by the General Research Fund of the Hong Kong Research Grants Council (RGC) [Grant No. 16204519]. S.F.A. was supported by the Hong Kong Ph.D. Fellowship Scheme (HKPFS).

---

**Figure S1.**
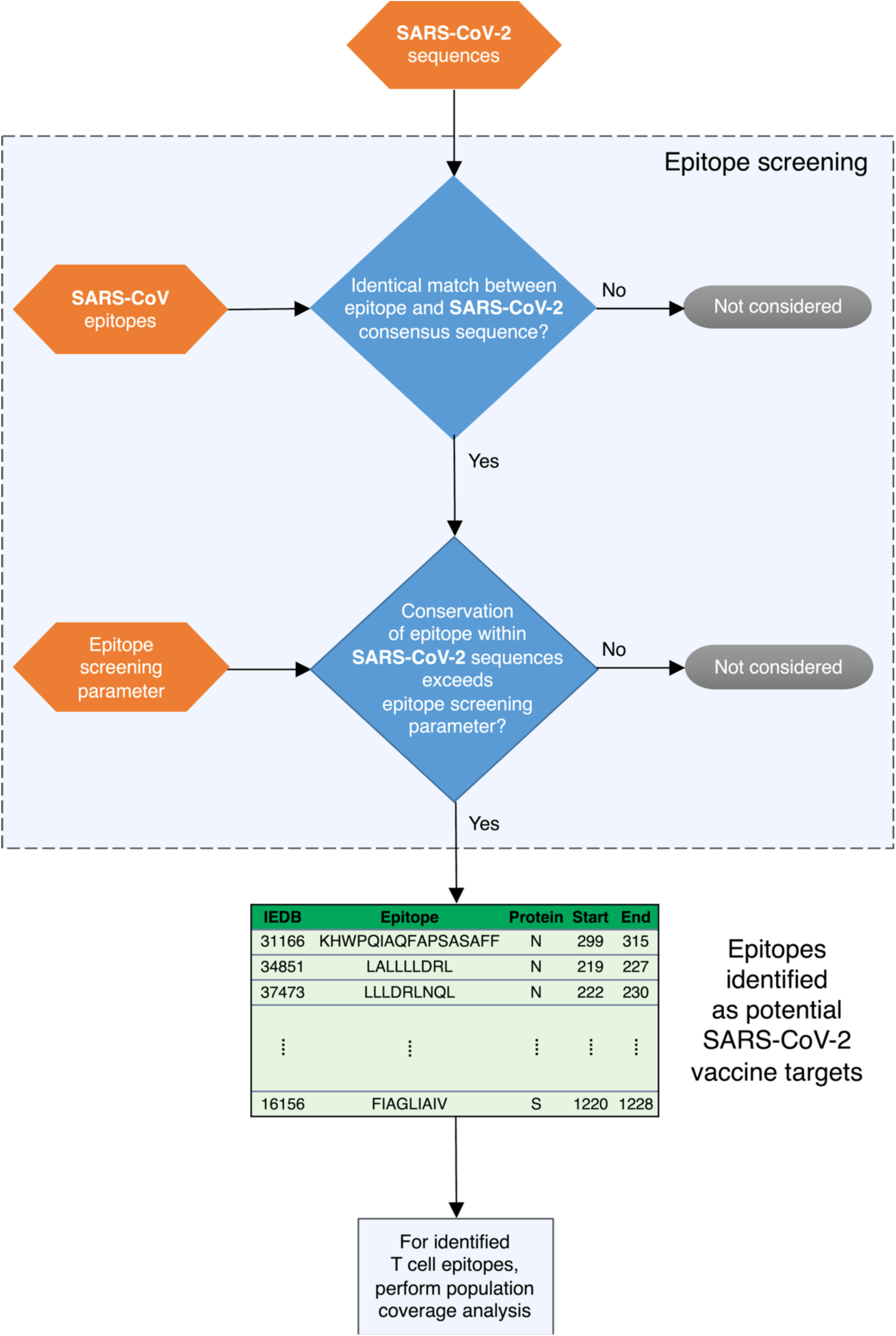
Epitope screening protocol used by COVIDep for providing vaccine target recommendations for SARS-CoV-2. COVIDep periodically pools SARS-CoV-2 sequence data and compares with experimentally-determined B cell and T cell epitopes of SARS-CoV. The system outputs those epitopes that are genetically similar in SARS-CoV-2, based on an epitope screening parameter. This user-defined parameter allows the user to select epitopes based on their conservation in the SARS-CoV-2 sequence data, where conservation is defined as the fraction of SARS-CoV-2 sequences with the exact epitope sequence. For the identified T cell epitopes, population coverage analysis is performed to estimate the percentage of a specified population that can elicit a response against them.

